# CRISPR-induced indels and base editing using the *Staphylococcus aureus* Cas9 in potato

**DOI:** 10.1101/2020.06.26.173138

**Authors:** Florian Veillet, Marie-Paule Kermarrec, Laura Chauvin, Jean-Eric Chauvin, Fabien Nogué

## Abstract

Genome editing is now widely used in plant science for both fundamental research and molecular crop breeding. The clustered regularly interspaced short palindromic repeats (CRISPR) technology, through its precision, high efficiency and versatility, allows to edit many sites in plant genomes. This system has been highly successful to produce of knock-out mutants through the introduction of frameshift mutations due to error-prone repair pathways. Nevertheless, recent new CRISPR-based technologies such as base editing and prime editing can generate precise and on request nucleotide conversion, allowing to fine-tune protein function and generate gain-of-function mutants. However, genome editing through CRISPR systems still have some drawbacks and limitations, such as the PAM restriction and the need for more diversity in CRISPR tools to simultaneously mediate different catalytic activities. In this study, we successfully used the CRISPR-Cas9 system from *Staphylococcus aureus* (SaCas9) for the introduction of frameshift mutations in the tetraploid genome of the cultivated potato (*Solanum tuberosum*). We also developed a *S. aureus*-cytosine base editor that mediate nucleotide conversions, allowing to precisely modify specific residues or regulatory elements in potato. Our proof-of-concept results in potato expand the plant dicot CRISPR toolbox for biotechnology and precision breeding applications.

## Introduction

The recent and considerable development of plant genome editing in the last few years has opened new avenues and exciting perspectives for both fundamental research and crop breeding. The class 2 type II CRISPR-Cas9 genome editing system from *Streptococcus pyogenes* has been broadly adopted by the plant science community, and consists of a two-components complex made of the DNA endonuclease SpCas9 and a customizable single guide RNA (sgRNA) [1]. This complex scans the genome in search of a 5’-NGG-3’ protospacer adjacent motif (PAM), triggering local DNA melting and interrogation of adjacent DNA sequence for complementarity with the customizable spacer sequence at the 5’end of the sgRNA, eventually resulting in double strand DNA break (DSB) about 3-bp upstream the PAM by the concerted activity of HNH and RuvC nuclease domains [2]. Once a DSB is created, the error-prone non-homologous end-joining (NHEJ) DNA repair pathway is activated [3], eventually resulting in unfaithful DNA repair that generate random small insertions or deletions (indels) mutations at the breaking site, typically leading to gene knockout through frameshift mutations.

While most studies focused on the production of loss-of-function alleles so far, new CRISPR tools have been recently developed, such as the CRISPR-mediated base editing system that allows precise base conversion without neither a donor DNA or the induction of a DSB [4]. So far, two kinds of base editors (BEs) have been developed: cytosine base editors (CBEs) [5] and adenine base editors (ABEs) [6] whose architecture is composed of the fusion of a Cas9 with an impaired DNA cleavage activity, mostly a nickase Cas9 (nCas9) for plant applications, and a catalytic domain mediating cytosine or adenine deamination, respectively. During the fixation of the nCas9 to its genomic target, a small window of the non-targeted ssDNA can serve as a substrate for deaminase domains. While ABEs almost exclusively mediate A-to-G conversion [6], CBEs can result in C-to-T, C-to-G and C-to-A according to the architecture of the BE [7].

Although the CRISPR-SpCas9 system revolutionized plant functional genomics, several other Cas9 enzymes from diverse bacteria have been used as an alternative for genome editing in plants, including the *Staphylococcus aureus* Cas9 (SaCas9) [8,9]. Use of SaCas9 for plant genome editing presents some assets. First, because the PAM recognized by the SaCas9 (5’-NNGRRT-3’) is different from the canonical 5’-NGG-3’ PAM from SpCas9 (where N is for any nucleotide while R can be A or G), its use expands the number of sites that can be targeted in a given genome. In addition, the fact that the PAM of SaCas9 is more sophisticated than the one from SpCas9 may allow to increase the specificity of the system by limiting the off-target activity, especially for highly conserved genomic regions that are frequent in polyploid species. Finally, because SaCas9 is smaller than SpCas9 (1053 vs 1368 amino acids), delivering into plant cells could be easier, especially for strategies involving virus vectors. To date, CRISPR-SaCas9 has been applied in different plant species for both gene knockout and/or base editing applications, including tobacco [10], *Arabidopsis [11]*, citrus [12] and rice [10,13,14].

The cultivated and tetraploid potato (*Solanum tuberosum*) received much attention for genome editing in the last few years by several groups. These achievements allowed to produce plants with new agronomic traits with gene knockout and/or base editing approaches, such as the production of tubers with low levels of amylose [15–17] or tubers with improved resistance to harvest and post-harvest procedures [18]. However, all the studies on potato used the classical or engineered variants [19] of SpCas9 so far, pointing out to the necessity to broaden the CRISPR toolbox for this species that constitutes one of the most important crops for food production worldwide. In this study, we report on the successful use of the SaCas9 enzyme for both knockout and base editing applications in the tetraploid potato, confirming that the CRISPR-SaCas9 system constitutes a relevant alternative to the classical CRISPR-SpCas9 technology for functional studies and plant breeding.

## Results and Discussion

### CRISPR-SaCas9-mediated gene editing of the potato genome

To evaluate the efficiency of the CRISPR-SaCas9 system in potato, we first designed two sets of two sgRNAs each, whose expression was driven by the *Arabidopsis* U6-26 promoter [11]. The first set targeted the *StGBSSI* and *StDMR6-1* genes with spacers of 20-bp sequence length (sgRNA1 and 2), while the second set targeted the same loci but with spacers of 24-bp sequence length (sgRNA3 and 4) (Fig 1). All the spacer sequences were chosen upstream of a 5’-NNGGAT-3’ PAM with a high specificity score according to the CRISPOR software, selecting spacer sequences harboring at least 4 mismatches with other loci in the genome. For expression of the CRISPR-SaCas9 system in potato cells, we cloned each set of sgRNA cassettes into the binary vector previously used in *Arabidopsis [11]*, resulting into the pDeSaCas9/sgRNA1-2 and the pDeSaCas9/sgRNA3-4 plasmids (Fig 1).

**Fig 1:**
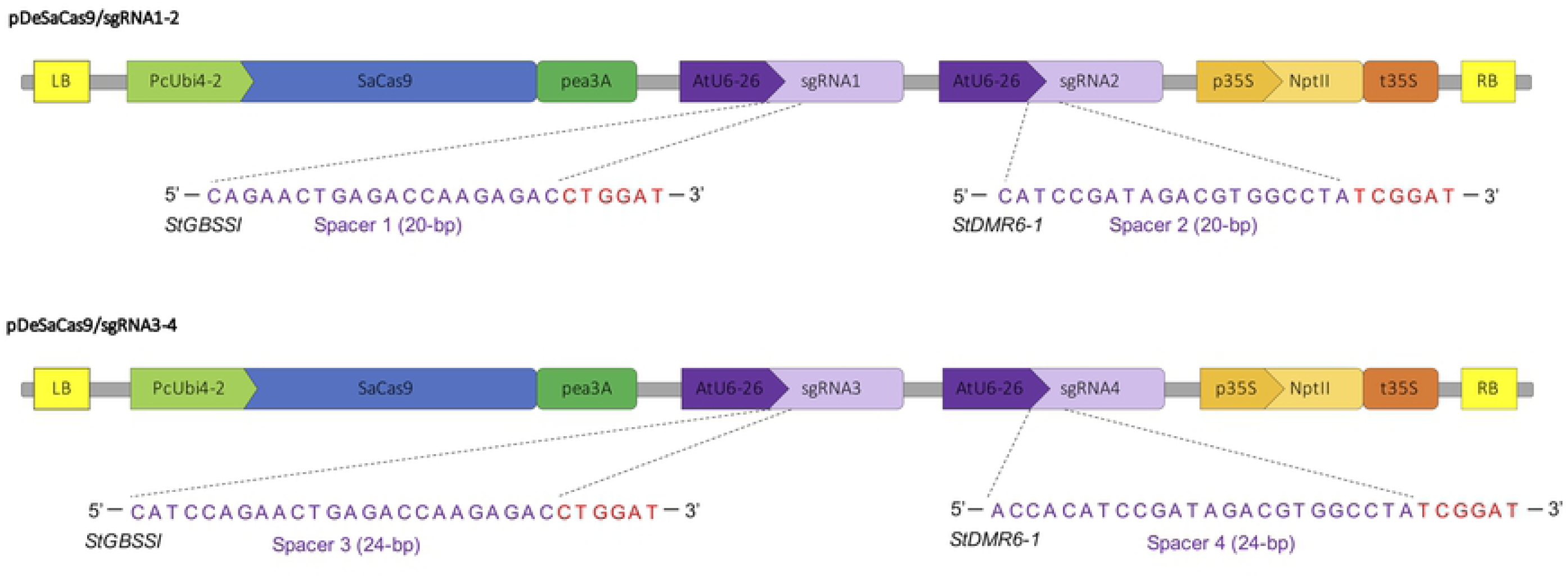
CRISPR-SaCas9 plasmids for genome editing in potato. Schematic representation of the two CRISPR-SaCas9 binary plasmids used for editing the *StGBSSI* and *StDMR6-1* targeted sites. For each sgRNA (indicated with a number from 1 to 4), the genomic targeted site is represented with the spacer and the PAM sequences in purple and red, respectively. LB: left border of T-DNA; RB: right border of T-DNA; PcUbi4-2: *Petroselinum crispum* Ubiquitin4-2 promoter; Pea3A: *Pisum sativum* 3A terminator, AtU6-26: *Arabidopsis* U6-26 promoter; p35S: CaMV 35S promoter; promoter; nptII: neomycin phosphotransferase; t35S: CaMV 35S terminator. The schemes are not at scale and are for illustrative purposes only.

To deliver the CRISPR components into potato cells, we performed an *Agrobacterium*-mediated transformation of potato explants, and genomic DNA from regenerative plants selected on kanamycin-supplemented medium was analyzed by high resolution melting (HRM) analysis followed by Sanger sequencing. For the pDeSaCas9/sgRNA1-2 condition, among the 33 plants that rooted on kanamycin-containing medium, none of them was mutated at the *StGBSSI* locus (sgRNA1), while 11 plants (33% efficiency) were found to harbor mutations in the *StDMR6-1* target sequence (sgRNA2) (Fig 2A). For the pDeSaCas9/sgRNA3-4 condition, among the 27 plants that developed roots on kanamycin-containing medium, none of them displayed mutations at the *StGBSSI* locus (sgRNA3), while 4 plants (15% efficiency) were found to be mutated at the *StDMR6-1* target site (sgRNA4) (Fig 2A). These results indicate that the SaCas9 can be used for gene editing in the potato genome, with spacer sequences of up to 24-bp. The observation that no editing activity was detected at the *StGBSSI* target locus for both spacer lengths (sgRNA1 and 3) may be due to the presence of an inefficient motif in the spacer sequence. However, none of the two motifs identified as inefficient in a previous study [20] was present in our spacer sequences, suggesting another origin for the lack of editing at this locus, such as the genomic context that may interfere with Cas9 binding and cleavage, and pointing out to the necessity to test independent spacer sequences for a target gene in order to maximize the likelihood of successful editing.

**Fig 2:**
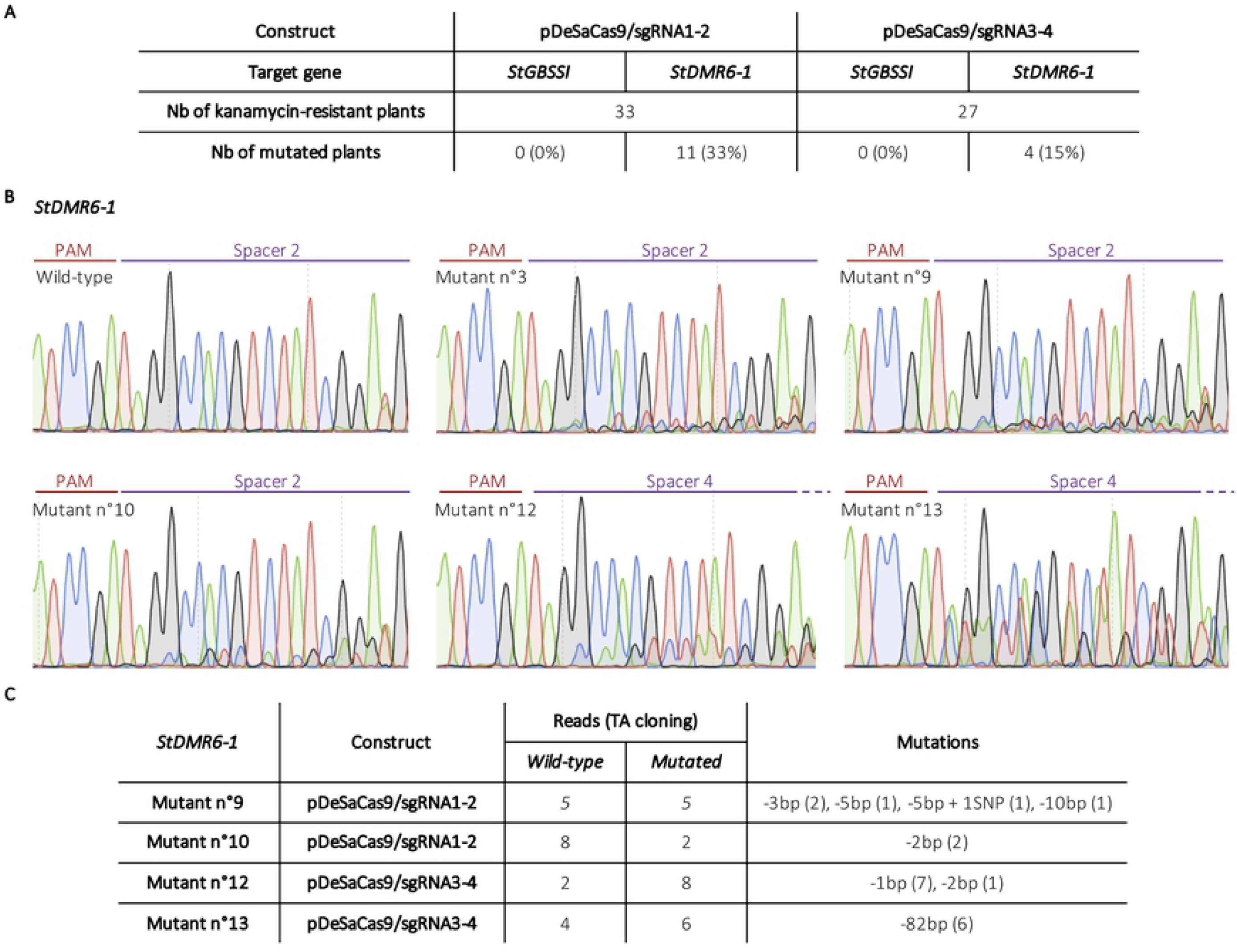
CRISPR-SaCas9-mediated genome editing in potato. **A)** Table summarizing the editing efficiencies at the *StGBSSI* and *StDMR6-1* targeted loci using both HRM analysis and Sanger sequencing. **B)** Sanger chromatograms of some CRISPR-SaCas9-edited potato plants at the *StDMR6-1* gene with the pDeSaCas9/sgRNA1-2 for mutants n°3, 9 and 10 and with the pDeSaCas9/sgRNA3-4 for mutants n°12 and 13. The PAM, which is located on the reverse strand, is indicated in red and the spacer sequence in purple. **C)** Table summarizing the results of Sanger sequencing for 4 mutants after cloning of individual PCR products through TA cloning. Mutants n°9 and 10 and mutants n°12 and 13 were edited using the pDeSaCas9/sgRNA1-2 and pDeSaCas9/sgRNA3-4 constructs, respectively.

Based on manual analysis of the chromatograms for sgRNA2 and 4 (*StDMR6-1* locus), we found that frameshift mutations mostly occurred about 4-bp upstream of the PAM sequence (Fig 2B), as previously reported for SaCas9 [11,12]. For each chromatogram, we found a clearly identifiable wild-type sequence trace (Fig 2B), indicating that CRISPR-induced indels did not occur for all the alleles. Sanger sequencing analysis of the *StDMR6-1* targeted sequence in the wild-type (Desiree cultivar) identified one SNP (T/A) (Fig 2B), that is present at the 5’end of target sequence of sgRNA2 and sgRNA4 (position -18 from the PAM), which is supposed to be present on two alleles according to a recently released SNP map from the Desiree genome [21]. The absence of mutants affected on the four-alleles may be due to the presence of this natural SNP, thereby affecting overall editing efficiency. To characterize in more details the SaCas9-mediated editing footprint at the *StDMR6-1* target site, we sequenced individual PCR amplicon after a TA-cloning reaction for 4 independent mutated plants. Most of the mutations were small indels about 3/4-bp upstream of the PAM (Fig 2C and S1 Fig 1), confirming the results from global PCR products sequencing. However, we also observed a 82-bp deletion for one plant, showing that large sequence rearrangement can occur at the target site (Fig 2C and S1 Fig 1). Intriguingly, we did not find any allele sequence harboring the natural SNP, suggesting that a bias occurred during the TA cloning reaction.

Taken together and compared to our previous work on genome editing in potato [17], our results show that SaCas9 constitutes an alternative to the classical SpCas9. As previous data showed that SaCas9 and SpCas9 could edit different plant genomes with a comparable efficiency [10–12], the efficiency of SaCas9 in potato needs to be further investigated by targeting several other loci before any conclusion on the relative efficiency of SaCas9 compared to SpCas9 in potato.

### CRISPR-SanCas9-mediated cytosine base editing of the potato genome

Because introducing precise nucleotide substitutions is of upmost importance for both functional genomics (e.g. protein domain characterization) and plant breeding (e.g. gain of function variants), and because knockout mutants can have growth penalties compared to functional allelic variants, we next sought to develop a CRISPR-SanCas9 cytosine base editor to mediate cytosine substitution. We first introduced a punctual mutation in the SaCas9 sequence to produce a SanCas9 (D10A) that we fused to a dicot codon-optimized fragment harboring both a cytosine deaminase (PmCDA1) and an uracil glycosylase inhibitor (UGI) domain. This fusion protein was then cloned into a modified version of the pDe backbone [19,22,23], resulting in the pDeSanCas9_PmCDA1_UGI binary plasmid for expression in dicot species (Fig 3A). The four sgRNAs used for CRISPR-mediated indels were individually cloned into this CBE through Gateway cloning (Fig 3B), each spacer harboring two to five cytosines in the putative editing window established for this CBE, based on previous studies using the PmCDA1 enzyme in plants with SpnCas9 or SanCas9 [13,17,19,24,25] (Fig 3B).

**Fig 3:**
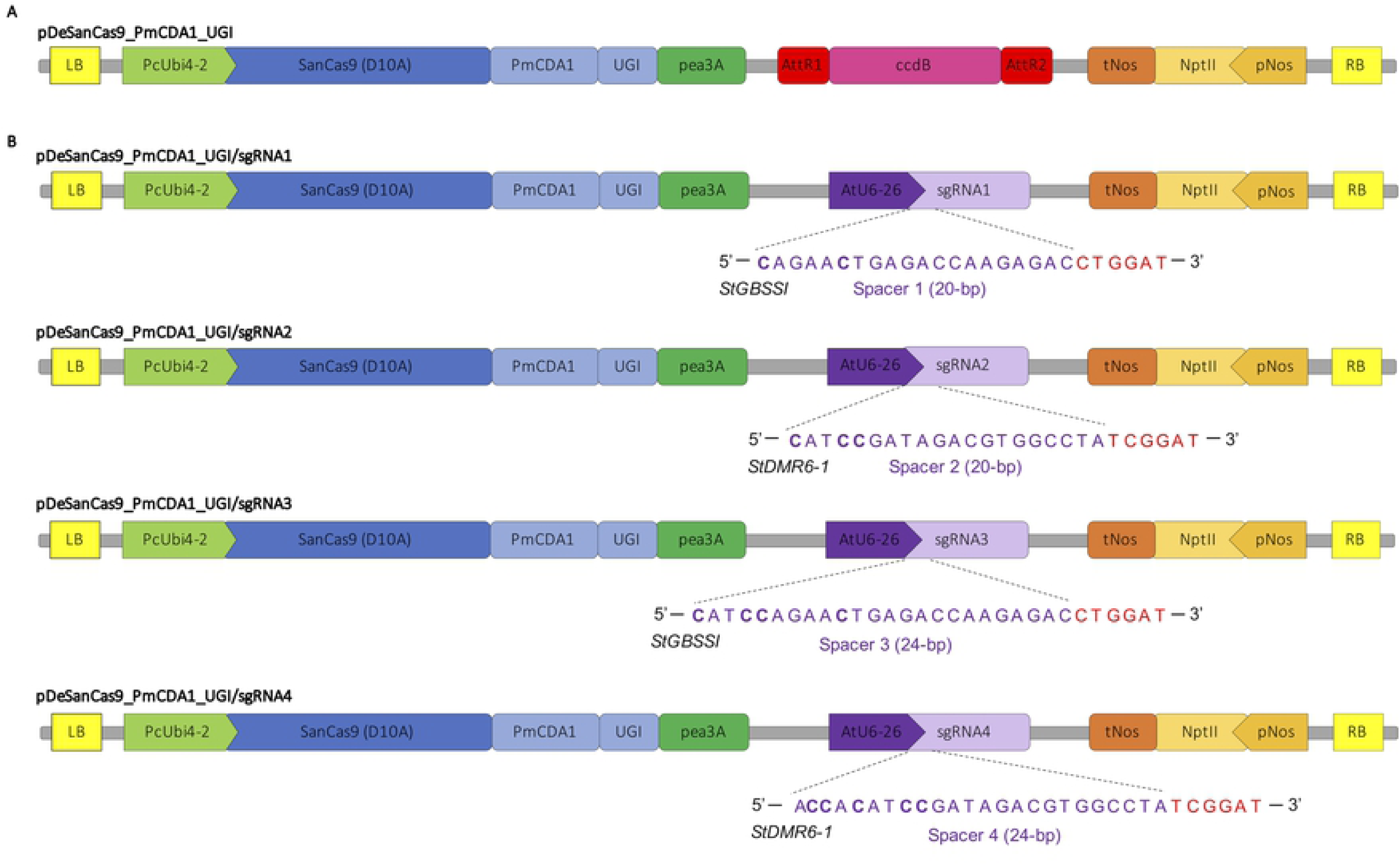
CRISPR-SaCBE plasmids for base editing in potato. **A)** Partial schematic representation of the CRISPR-SaCBE binary plasmid produced for expression in dicot species. This empty destination plasmid allows for the introduction of guide cassette through Gateway LR reaction. **B)** Partial schematic representation of the four CRISPR-SaCBE binary plasmids used for base editing at the *StGBSSI* and *StDMR6-1* targeted sites. For each sgRNA (indicated with a number from 1 to 4), the genomic targeted site is represented with the spacer and the PAM sequences in purple and red, respectively. The cytosines that are located in the putative edition window of the CBE are represented in bold. AttR1 and AttR2 corresponds to the Gateway cloning recombination sequences for the cloning of the guide cassette; LB: left border of T-DNA; RB: right border of T-DNA; PcUbi4-2: Petroselinum *crispum* Ubiquitin4-2 promoter; PmCDA1: *Petromyzon marinus* cytidine deaminase; UGI: uracil glycosylase inhibitor; Pea3A: *Pisum sativum* 3A terminator, AtU6-26: *Arabidopsis* U6-26 promoter; pNos: nopaline synthase promoter; nptII: neomycin phosphotransferase; tNos: nopaline synthase terminator. The schemes are not at scale and are for illustrative purposes only.

The delivery of the CBEs into potato cells was performed through *Agrobacterium*-mediated transformation and potato explants were then grown on kanamycin-containing medium for several weeks. For both constructs targeting the *StGBSSI* gene (sgRNA1 and sgRNA3), none of the regenerated plants displayed mutations according to HRM analysis, which indicates, together with the inability to induce indels at this locus with the SaCas9 nuclease, that the spacer sequences and/or the targeted locus display characteristics preventing an efficient fixation of the CRISPR complex. For the pDeSanCas9_PmCDA1_UGI/sgRNA2 construct harboring a 20-bp spacer sequence, we did not find any base edited plant, suggesting that cytosine deamination occur with lower efficiency than dsDNA cleavage at this locus. However, we identified three mutated plants for the pDeSanCas9_PmCDA1_UGI/sgRNA4 construct that harbors a 24-bp spacer sequence. One of these mutants (#16) experienced indel mutations at one or more targeted alleles, while two mutants (#17 and #18) correspond to cleanly base edited plants (Fig 4). Although this 24-bp spacer sequence was less efficient than the corresponding 20-bp spacer sequence for inducing indels mutations (Fig 2A), its higher efficiency for cytosine base editing may be due to the presence of additional cytosines in the editing window at the 5’end of the spacer (Fig 3B). Supporting this hypothesis, we found that base conversion only occurred at C_-23_ and C_-22_ (counting from the PAM) in the two cleanly base edited plants (Fig 4B). Interestingly, despite the presence of one UGI domain, our CBE construct was able to mediate both transition (C-to-T) and transversion (C-to-G) mutations (Fig 4), allowing to diversify the edits, albeit at the cost of indel formation that occurred at the 5’end of the target sequence for one plant. This observation suggests that cytosine deamination-associated DNA repair mechanisms are involved in the production of this by-product.

**Fig 4:**
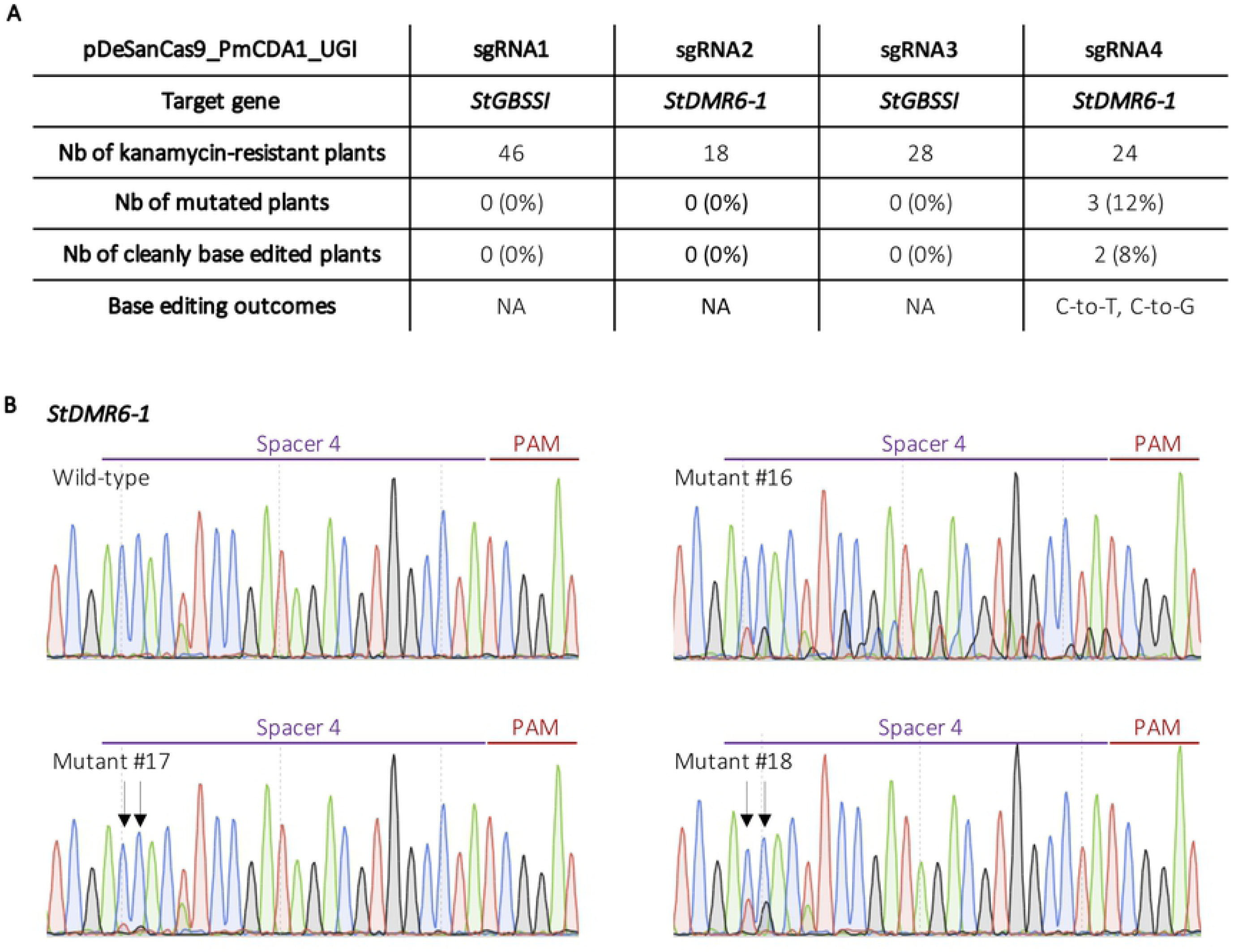
CRISPR-SaCBE-mediated base editing in potato. **A)** Table summarizing the base editing efficiencies at the *StGBSSI* and *StDMR6-1* targeted loci using both HRM analysis and Sanger sequencing. Cleanly base edited plants refers to plants that harboured cytosine conversion without the introduction on indels in the target sequence. **B)** Sanger chromatograms of the three CRISPR-SaCBE-edited potato plants at the *StDMR6-1* gene with the pDeSanCas9_PmCDA1_UGI/sgRNA4. Because the PAM (in red) is located on the reverse strand, and in order to avoid confusion, we sequenced using a reverse primer to clearly identify the C conversion. The spacer sequence is represented in purple.

To summarize, the CRISPR-SanCas9 CBE was able to achieve cytosine base conversion at distal location from the 5’-NNGGAT-3’ PAM in the cultivated potato, which is to our knowledge the first report of such application in a dicot species. The CRISPR-SanCas9 CBE developed for this study represent a complementary tool to the previously described SpnCas9 based CBE and, thanks to its capacity to hybridize with 18-24-bp guide sequences [26], may be useful to efficiently target specific nucleotide at the distal part of longer spacer sequence, as demonstrated here.

## Concluding remarks

The CRISPR-SaCas9 tools used and developed in this study broaden the scope of genome editing applications for potato, but also for dicot species in general. While the use of SaCas9 that recognizes a sophisticated 5’-NNGRRT-3’ PAM may be useful to limit off-target activity at conserved sequences, this enzyme suffers from a narrowed targeting scope for base editing experiments due to the low occurrence of the PAM and the necessity to place the targeted base(s) in small edition window. Therefore, to unleash the base editing potential of SanCas9, the SanCas9-KKH engineered variant that recognizes the relaxed 5’-NNNRRT-3’ PAM has been successfully used in rice for both adenine and cytosine conversion [13,14,27], and could be of particular interest in dicot species. Finally, validation in potato of the use of SpCas9 and SaCas9, that associate with distinct sgRNA scaffolds, makes possible their simultaneous use to perform different catalytic functions (e.g. gene knock out, base editing, prime editing, transcription regulation, epigenome modulation) in a single transformation step and extent the possibilities of genome engineering in this essential crop.

## Material and Methods

### Plant material

The potato cultivar Desiree (ZPC, the Netherlands) was propagated in sterile conditions in 1X MS medium including vitamins at pH 5.8 (Duchefa, the Netherlands), 0.4 mg/L thiamine hydrochloride (Sigma-Aldrich, USA), 2.5% sucrose and 0.8% agar powder (VWR, USA). Plants were cultured *in vitro* in a growth chamber at 19°C with a 16:8 h L/D photoperiod.

### Cloning procedures

The entry plasmid pEn_Sa_Chimera for spacer cloning and the binary vector pDeSaCas9 were kindly provided by Holger Puchta [11]. For spacer cloning, the pEn_Sa_Chimera entry plasmid was digested by *Bbs*I and annealed oligonucleotides bearing complementary overhangs were ligated through T4 DNA ligase (ThermoFisher Scientific, USA) (S2 Table 1). For multiplex editing using the pDeSaCas9/sgRNA1-2 and pDeSaCas9/sgRNA3-4, sgRNA1 and sgRNA3 were introduced into the pDeSaCas9 backbone through *Mlu*I restriction and T4 DNA ligation (ThermoFisher Scientific, USA), while sgRNA2 and sgRNA4 were then introduced through a LR Gateway reaction (ThermoFisher Scientific, USA). The resulting plasmids were checked by restriction digestion and Sanger sequencing (Fig 1 and S2 Table 1).

The pDeSanCas9_PmCDA1_UGI binary plasmid was produced as follow. The SanCas9 sequence was produced through PCR amplification with the Superfi DNA polymerase (ThermoFisher Scientific, USA) using a forward primer bearing polymorphism for D10A amino acid shift (S2 Table 1), devoid of a STOP codon. The PCR fragment was cloned into an intermediate pTwist plasmid through *Mlu*I/*Eco*RI restriction followed by T4 DNA ligation (ThermoFisher Scientific, USA). A sequence encoding the PmCDA1 and UGI catalytic domains was previously dicot-codon optimized and synthesized (TwistBioscience, USA) [19], and cloned into the intermediate pTwist plasmid through *EcoR*I restriction and T4 DNA ligation (ThermoFisher Scientific, USA), downstream of the SanCas9 coding sequence. The construct was checked by sanger sequencing (S2 Table 1). The SanCas9_PmCDA1_UGI sequence (S1 Fig 2) was then cloned into a modified pDeCas9 backbone [23] through *Asc*I restriction and T4 DNA ligation (ThermoFisher Scientific, USA). The final pDeSanCas9_PmCDA1_UGI was checked by restriction ligation and Sanger sequencing (Fig 3A and S2 Table 1). Previously built sgRNA cassettes were then individually cloned into the Sa_CBE plasmid through a LR Gateway reaction (ThermoFisher Scientific, USA). The resulting plasmids were checked by restriction digestion and Sanger sequencing (Fig 3B and S2 Table 1).

### *Agrobacterium*-mediated transformation and plant regeneration

Binary plasmids described above were transferred into *Agrobacterium* C58pMP90 strain by heat shock. *Agrobacterium*-mediated stable plant transformation and plant regeneration were performed on explants of the Desiree cultivar, as previously described [17]. Plant tissues were cultured on 50 mg/L kanamycin, and regenerated plants were then transferred to a culture medium containing 50 mg/L kanamycin or tested for the presence of the T-DNA by PCR (S2 Table 1).

### Target site genotyping

Genomic DNA from control and regenerated plants was extracted using the NucleoSpin Plant II kit (Macherey–Nagel, Germany) according to the manufacturer’s instructions. HRM analysis was performed using the High Resolution Melting Master (Roche Applied Science, Germany) on the LightCycler® 480 II system (Roche Applied Science, Germany) (S2 Table 1), as previously described [17]. Plants harboring a HRM mutated profile were then Sanger sequenced (Genoscreen, France) (S2 Table 1). Some plants harboring mutations at the *StDMR6-1* locus with the pDeSaCas9 constructs were further analyzed by cloning the PCR products (Superfi DNA polymerase, ThermoFisher Scientific, USA) into the pCR4-TOPO TA vector (ThermoFisher Scientific, USA), followed by Sanger sequencing (Genoscreen, France).

## Acknowledgments

We thank Holger Puchta and his team (Botanical Institute II, Karlsruhe Institute of Technology, Karlsruhe, Germany) for providing the pDeSaCas9 backbone. We thank Peter Rogowsky for his efficient management of the GENIUS project. We acknowledge the BrACySol BRC (INRAE, Ploudaniel, France) for providing the potato plants used in this study.

## Supporting information

**S1 Fig1: Alignment of Sanger chromatograms obtained after TA cloning of individual PCR fragments of the StDMR6-1 targeted locus**. The reference sequence is displayed at the top of each panel, with the position of the PAM (in red, on the reverse strand) and the spacer sequences (in blue). The number on the right of each chromatogram corresponds to the number of identical chromatograms observed. The Geneious software was used for the alignments.

**S1 Fig 2: Coding sequence of the Sa-CBE developed in this study**. The SanCas9 sequence is in blue, the two NLS sequences in purple, the PmCDA1 sequence in green and the UGI sequence in red. All the coding sequence is optimized for expression in dicot species.

**S2 Table 1: List of primers used in this study**.

## References

1. Jinek M, Chylinski K, Fonfara I, Hauer M, Doudna JA, Charpentier E. A Programmable Dual-RNA-Guided DNA Endonuclease in Adaptive Bacterial Immunity. Science. 2012;337: 816–821. doi: 10.1126/science.1225829

2. Jiang F, Doudna JA. CRISPR–Cas9 Structures and Mechanisms. Annu Rev Biophys. 2017;46: 505–529. doi: 10.1146/annurev-biophys-062215-010822

3. Puchta H. The repair of double-strand breaks in plants: mechanisms and consequences for genome evolution. Journal of Experimental Botany. 2004 [cited 7 Jun 2020]. doi: 10.1093/jxb/eri025

4. Mishra R, Joshi RK, Zhao K. Base editing in crops: current advances, limitations and future implications. Plant Biotechnol J. 2020;18: 20–31. doi: 10.1111/pbi.13225

5. Nishida K, Arazoe T, Yachie N, Banno S, Kakimoto M, Tabata M, et al. Targeted nucleotide editing using hybrid prokaryotic and vertebrate adaptive immune systems. Science. 2016;353: aaf8729–aaf8729. doi: 10.1126/science.aaf8729

6. Gaudelli NM, Komor AC, Rees HA, Packer MS, Badran AH, Bryson DI, et al. Programmable base editing of A•T to G•C in genomic DNA without DNA cleavage. Nature. 2017;551: 464–471. doi: 10.1038/nature24644

7. Komor AC, Zhao KT, Packer MS, Gaudelli NM, Waterbury AL, Koblan LW, et al. Improved base excision repair inhibition and bacteriophage Mu Gam protein yields C:G-to-T:A base editors with higher efficiency and product purity. Sci Adv. 2017;3: eaao4774. doi: 10.1126/sciadv.aao4774

8. Zhang Y, Malzahn AA, Sretenovic S, Qi Y. The emerging and uncultivated potential of CRISPR technology in plant science. Nat Plants. 2019;5: 778–794. doi: 10.1038/s41477-019-0461-5

9. Murovec J, Pirc Ž, Yang B. New variants of CRISPR RNA-guided genome editing enzymes. Plant Biotechnol J. 2017;15: 917–926. doi: 10.1111/pbi.127362915

10. Kaya H, Mikami M, Endo A, Endo M, Toki S. Highly specific targeted mutagenesis in plants using Staphylococcus aureus Cas9. Sci Rep. 2016;6: 26871. doi: 10.1038/srep26871

11. Steinert J, Schiml S, Fauser F, Puchta H. Highly efficient heritable plant genome engineering using Cas9 orthologues from Streptococcus thermophilus and Staphylococcus aureus. Plant J. 2015;84: 1295–1305. doi: 10.1111/tpj.13078

12. Jia H, Xu J, Orbović V, Zhang Y, Wang N. Editing Citrus Genome via SaCas9/sgRNA System. Front Plant Sci. 2017;8: 2135. doi: 10.3389/fpls.2017.02135

13. Qin R, Li J, Li H, Zhang Y, Liu X, Miao Y, et al. Developing a highly efficient and wildly adaptive CRISPR-SaCas9 toolset for plant genome editing. Plant Biotechnol J. 2019;17: 706–708. doi: 10.1111/pbi.13047

14. Hua K, Tao X, Zhu J-K. Expanding the base editing scope in rice by using Cas9 variants. Plant Biotechnol J. 2019;17: 499–504. doi: 10.1111/pbi.12993

15. Andersson M, Turesson H, Nicolia A, Fält A-S, Samuelsson M, Hofvander P. Efficient targeted multiallelic mutagenesis in tetraploid potato (Solanum tuberosum) by transient CRISPR-Cas9 expression in protoplasts. Plant Cell Rep. 2017;36: 117–128. doi: 10.1007/s00299-016-2062-3

16. Andersson M, Turesson H, Olsson N, Fält A-S, Ohlsson P, Gonzalez MN, et al. Genome editing in potato via CRISPR-Cas9 ribonucleoprotein delivery. Physiol Plantarum. 2018;164: 378–384. doi: 10.1111/ppl.12731

17. Veillet F, Chauvin L, Kermarrec M-P, Sevestre F, Merrer M, Terret Z, et al. The Solanum tuberosum GBSSI gene: a target for assessing gene and base editing in tetraploid potato. Plant Cell Rep. 2019;38: 1065–1080. doi: 10.1007/s00299-019-02426-w

18. González MN, Massa GA, Andersson M, Turesson H, Olsson N, Fält A-S, et al. Reduced Enzymatic Browning in Potato Tubers by Specific Editing of a Polyphenol Oxidase Gene via Ribonucleoprotein Complexes Delivery of the CRISPR/Cas9 System. Front Plant Sci. 2020;10: 1649. doi: 10.3389/fpls.2019.01649

19. Veillet F, Perrot L, Guyon-Debast A, Kermarrec M-P, Chauvin L, Chauvin J-E, et al. Expanding the CRISPR Toolbox in P. patens Using SpCas9-NG Variant and Application for Gene and Base Editing in Solanaceae Crops. IJMS. 2020;21: 1024. doi: 10.3390/ijms21031024

20. Graf R, Li X, Chu VT, Rajewsky K. sgRNA Sequence Motifs Blocking Efficient CRISPR/Cas9-Mediated Gene Editing. Cell Reports. 2019;26: 1098-1103.e3. doi: 10.1016/j.celrep.2019.01.024

21. Sevestre F, Facon M, Wattebled F, Szydlowski N. Facilitating gene editing in potato: a Single-Nucleotide Polymorphism (SNP) map of the Solanum tuberosum L. cv. Desiree genome. Sci Rep. 2020;10: 2045. doi: 10.1038/s41598-020-58985-6

22. Fauser F, Schiml S, Puchta H. Both CRISPR/Cas-based nucleases and nickases can be used efficiently for genome engineering in *Arabidopsis thaliana*. Plant J. 2014;79: 348–359. doi: 10.1111/tpj.12554

23. Danilo B, Perrot L, Mara K, Botton E, Nogué F, Mazier M. Efficient and transgene-free gene targeting using Agrobacterium-mediated delivery of the CRISPR/Cas9 system in tomato. Plant Cell Rep. 2019;38: 459–462. doi: 10.1007/s00299-019-02373-6

24. Shimatani Z, Kashojiya S, Takayama M, Terada R, Arazoe T, Ishii H, et al. Targeted base editing in rice and tomato using a CRISPR-Cas9 cytidine deaminase fusion. Nat Biotechnol. 2017;35: 441–443. doi: 10.1038/nbt.3833

25. Veillet F, Perrot L, Chauvin L, Kermarrec M-P, Guyon-Debast A, Chauvin J-E, et al. Transgene-Free Genome Editing in Tomato and Potato Plants Using Agrobacterium-Mediated Delivery of a CRISPR/Cas9 Cytidine Base Editor. IJMS. 2019;20: 402. doi: 10.3390/ijms20020402

26. Friedland AE, Baral R, Singhal P, Loveluck K, Shen S, Sanchez M, et al. Characterization of Staphylococcus aureus Cas9: a smaller Cas9 for all-in-one adeno-associated virus delivery and paired nickase applications. Genome Biol. 2015;16: 257. doi: 10.1186/s13059-015-0817-8

27. Zhang C, Wang F, Zhao S, Kang G, Song J, Li L, et al. Highly efficient CRISPR-SaKKH tools for plant multiplex cytosine base editing. The Crop Journal. 2020;8: 418–423. doi: 10.1016/j.cj.2020.03.002

